# Severe asthma is associated with a remodeling of the pulmonary arteries in horses

**DOI:** 10.1101/2020.04.15.042903

**Authors:** Serena Ceriotti, Michela Bullone, Mathilde Leclere, Francesco Ferrucci, Jean-Pierre Lavoie

## Abstract

Pulmonary hypertension and *cor pulmonale* are complications of severe equine asthma, as a consequence of pulmonary hypoxic vasoconstriction. However, as pulmonary hypertension is only partially reversible by oxygen administration, other etiological factors are likely involved. In human chronic obstructive pulmonary disease, pulmonary artery remodeling contributes to the development of pulmonary hypertension. In rodent models, pulmonary vascular remodeling is present as a consequence of allergic airway inflammation. The present study investigated the presence of remodeling of the pulmonary arteries in severe equine asthma, its distribution throughout the lungs, and its reversibility following long-term antigen avoidance strategies and inhaled corticosteroid administration. Using histomorphometry, the total wall area of pulmonary arteries from different regions of the lung of asthmatic horses and controls was measured. The smooth muscle mass of pulmonary arteries was also estimated on lung sections stained for α-smooth muscle actin. Reversibility of vascular changes in asthmatic horses was assessed after 1 year of antigen avoidance alone or treatment with inhaled fluticasone. Pulmonary arteries showed increased wall area in apical and caudodorsal lung regions of asthmatic horses in both exacerbation and remission. The pulmonary arteries smooth muscle mass was similarly increased. Both treatments reversed the increase in wall area. However, normalization of the vascular smooth muscle mass was observed only after treatment with antigen avoidance, but not with fluticasone. In conclusion, severe equine asthma is associated with remodeling of the pulmonary arteries consisting in an increased smooth muscle mass. The resulting narrowing of the artery lumen could enhance hypoxic vasoconstriction, contributing to pulmonary hypertension. Vascular smooth muscle mass normalization is better achieved by antigen avoidance than with inhaled corticosteroids.

## Introduction

Severe equine asthma (also known as Heaves and Recurrent Airway Obstruction) is a chronic, non-infectious inflammatory lower airway disease, characterized by recurrent episodes of airway obstruction induced by exposure to environmental antigens (including fungi, hay mites and endotoxins) (2-4). Pulmonary hypertension and reversible *cor pulmonale* are reported as clinical complications of severe equine asthma (5-7). Diffuse airway obstruction increases regions of alveolar dead space. The resulting ventilation/perfusion mismatch impairs normal pulmonary gas exchanges, leading to alveolar hypoxia and eventually hypoxemia (8, 9). Chronic hypoxia and hypoxemia exacerbate sustained hypoxic pulmonary vasoconstriction, increasing pulmonary vascular resistance and contribute to pulmonary hypertension onset and progression (10). In asthmatic horses, airway obstruction was recently shown to induce pulmonary hypertension with right ventricular structural and functional alterations (11). Their pulmonary artery pressure, measured by invasive right heart catheterization, inversely correlates with arterial oxygen tension, suggesting a significant role of hypoxic pulmonary vasoconstriction in increasing pulmonary vascular resistance and subsequently the mean pulmonary artery pressure (11, 12). However, pulmonary hypertension is only partially reversed by oxygen administration and asthmatic horses have a significantly higher pulmonary artery pressure compared to healthy horses, even during disease remission (11, 12). Therefore, other factors likely contribute to pulmonary hypertension development onset in asthmatic horses.

In experimental rodent models, chronic hypoxia, hypoxemia and allergic airway inflammation induce pulmonary vascular remodeling that mainly involves small muscular pulmonary arteries (13-19). Pulmonary vascular remodeling is also well documented in chronic inflammatory respiratory disorders in humans such as chronic obstructive pulmonary disease (COPD) where it is recognized as a determining factor in the onset of pulmonary hypertension and *cor pulmonale* (20, 21). Remodeled muscular arteries show increased smooth muscle mass, resulting in artery wall thickening, lumen narrowing and vasoreactivity enhancement, with subsequent pulmonary vascular resistance augmentation. In mouse models of asthma, the increase in pulmonary arteries smooth muscle (PASM) mass also persists during short-term disease remission (13, 14).

Because hypoxemia and chronic airway inflammation also occur in severe equine asthma, they may induce remodeling of small pulmonary arteries. If present, pulmonary artery remodeling could then contribute to both pulmonary hypertension onset during clinical exacerbation of the disease and persistence of higher pulmonary artery pressure values during the disease remission. We therefore postulated that pulmonary vascular remodeling occurs in severe equine asthma and is associated with increased PASM mass and vascular lumen narrowing. According to our hypothesis, the present study had the following objectives: to 1) determine the presence and anatomical location of pulmonary artery remodeling in severe equine asthma; 2) assess whether differences are detectable between remission and exacerbation; 3) investigate the remodeling pattern and whether an increased PASM mass is present and; 4) assess remodeling reversibility after long-term asthma treatments.

## Materials and methods

### Study design and sample selection

This study was part of a larger project aiming to assess lung tissue remodeling and its reversibility in severe equine asthma. The present study focused on remodeling of pulmonary vasculature and particularly of muscular pulmonary arteries, namely small ramifications of the pulmonary arterial trunk that appear to be more prone to remodeling in previous studies on rodent models (18, 19). A blinded histomorphometric assessment was conducted on lung samples retrieved from the BTRE equine respiratory tissue biobank (http://www.btre.ca) in three different phases (study 1, 2 and 3). Study 1 consisted of a preliminary study performed on a subset of banked samples selected from a previous cadaveric study (1). Study 1 aimed to determine the presence of pulmonary artery remodeling and its anatomical location within different lung regions, as well as to assess differences between remission and exacerbation states. The lung samples were collected *post-mortem*, respectively from 6 asthmatic horses in exacerbation (antigen exposure), 6 asthmatic horses in remission (short-term antigen avoidance) and 6 age-matched controls. Samples belonged to different regions of the left lung and named cranio-caudally from “A” to “D” as previously described (1). Sample “E” was collected from the peripheral caudodorsal part of the main lung lobe and corresponds to the specific biopsy site during thoracoscopy. The wall area of muscular pulmonary arteries was measured by histomorphometry, as an estimation of the degree of artery wall thickening. Measurements were compared between asthmatic horses and controls in different lung regions. A comparison between asthmatic horses in remission and exacerbation was also performed.

Study 2 was performed on banked samples from a previous *in vivo* case-control study (22). Study 2 aimed to confirm *post-mortem* results and to determine the presence of PASM mass involvement in the remodeling. All samples belonged to peripheral lung biopsies collected via thoracoscopy from the caudodorsal lung of 6 asthmatic horses in remission and 5 age-matched controls, as previously described (23). The pulmonary artery wall area was measured, and the intimal and medial areas were evaluated, to assess presence of specific intimal and/or medial thickening. The PASM mass and the extracellular matrix (ECM) were quantified on sections immunostained for smooth muscle specific α-actin (α-SMA). Measurements were compared between asthmatic horses and controls.

Study 3 was performed on banked samples from an *in-vivo* prospective randomized clinical trial involving 11 asthmatic horses randomly divided into 2 groups and treated for 12 months with two different protocols (24). Study 3 aimed to assess reversibility of wall and PASM mass changes after long-term asthma treatment. Details concerning the animals used, treatment protocols and sampling procedures are reported elsewhere (24). Briefly, treatment protocols included antigen avoidance strategy alone for the whole experimental period (samples from 5 horses) or inhaled corticosteroid (fluticasone) alone for the first 6 months, combined with antigen avoidance for the remaining 6 months (samples from 6 horses). The pulmonary artery wall area and PASM mass were quantified by histomorphometry and compared in each treatment group between samples collected at baseline (before treatment, during disease exacerbation) and samples collected at follow-up (end of the treatment period).

### Sample processing and staining

Lung samples were fixed for 24 hours in 4% formaldehyde, embedded in paraffin, cut, placed on glass slides and stained. In study 1, samples were stained for histology with hematoxylin eosin saffron. In study 2 and 3, samples were stained for histology with modified Movat Russell pentachrome (25). This staining highlights the internal elastic lamina, allowing differentiation between intimal and medial layers within the artery wall. Samples for study 2 and 3 were also immunostained for α-SMA, as previously described (26).

### Histomorphometry

*Post-mortem* and *in viv*o histological sections were scanned and digitalized at 40X magnification as TIFF images with PanOptiq™ software version 1.4.3 (ViewsIQ, Vancouver, Canada); morphometric measurements were collected on TIFF images using ImageJ software version 1.4g (27). All the arteries with clear distinction between the wall and the *lumen* were evaluated. Both linear (1D) and bidimensional (2D) parameters were measured or derived, as an adaptation of what previously reported for human pulmonary vascular histomorphometry (28-30). Definition and collection methods of 1D and 2D “measured parameters” are summarized in figure 1 and table 1. “Derived parameters” were calculated from “measured parameters”, according to definitions summarized in table 2 (Office Excel^R^ software v. 2016; Microsoft Corporation, Redmond, USA). In horses, muscular pulmonary arteries have a minimal external diameter (ED) of 40 µm therefore, arteries with lower ED length were excluded (31). Narrowing index and GLA (great longitudinal axis)/ED *ratio* were used to assess differences due to histologic processing, artery collapse and cut sections respectively. Longitudinal cut sections of vessels (GLA/ED *ratio* > 3) were excluded, as previously suggested (30). The internal perimeter (IP) and the internal area (IA) were measured on Movat Russell pentachrome stained sections (study 2), only if the internal elastic lamina appeared well defined in the whole artery. The intimal area and the medial area were obtained only if IA measurement was available. To allow comparison among data of different sized-artery sections, these measurements were also expressed as percentages of the total area (wall area %, intimal area % and medial area %).

**Table 1:** 1D and 2D “measured parameters”. These parameters were adapted from those already defined in pulmonary vascular histomorphometric studies in human chronic obstructive pulmonary disease (COPD) (30). Those parameters marked with ^§^ were assessed only on sections stained with Movat Russell pentachrome (study 2).

| **1D measured parameters ($\mu\text{m}$)** |  |
| --- | --- |
| External artery diameter (ED) | Widest distance between external elastic laminae, perpendicular to the greatest longitudinal axis |
| Great longitudinal axis (GLA) | Longest distance between external elastic laminae |
| External perimeter (EP) | Outline of the external elastic lamina |
| Internal perimeter (IP) <sup>§</sup> | Outline of the internal elastic lamina |
| *Lumen* perimeter (LP) | Outline of the inner aspect of the intima |
| **2D measured parameters ($\mu\text{m}^2$)** |  |
| Total area (TA) | Area encompassed by the external perimeter |
| Internal area (IA) <sup>§</sup> | Area encompassed by the internal perimeter |
| *Lumen* area (LA) | Area encompassed by the lumen perimeter |

**Table 2:**
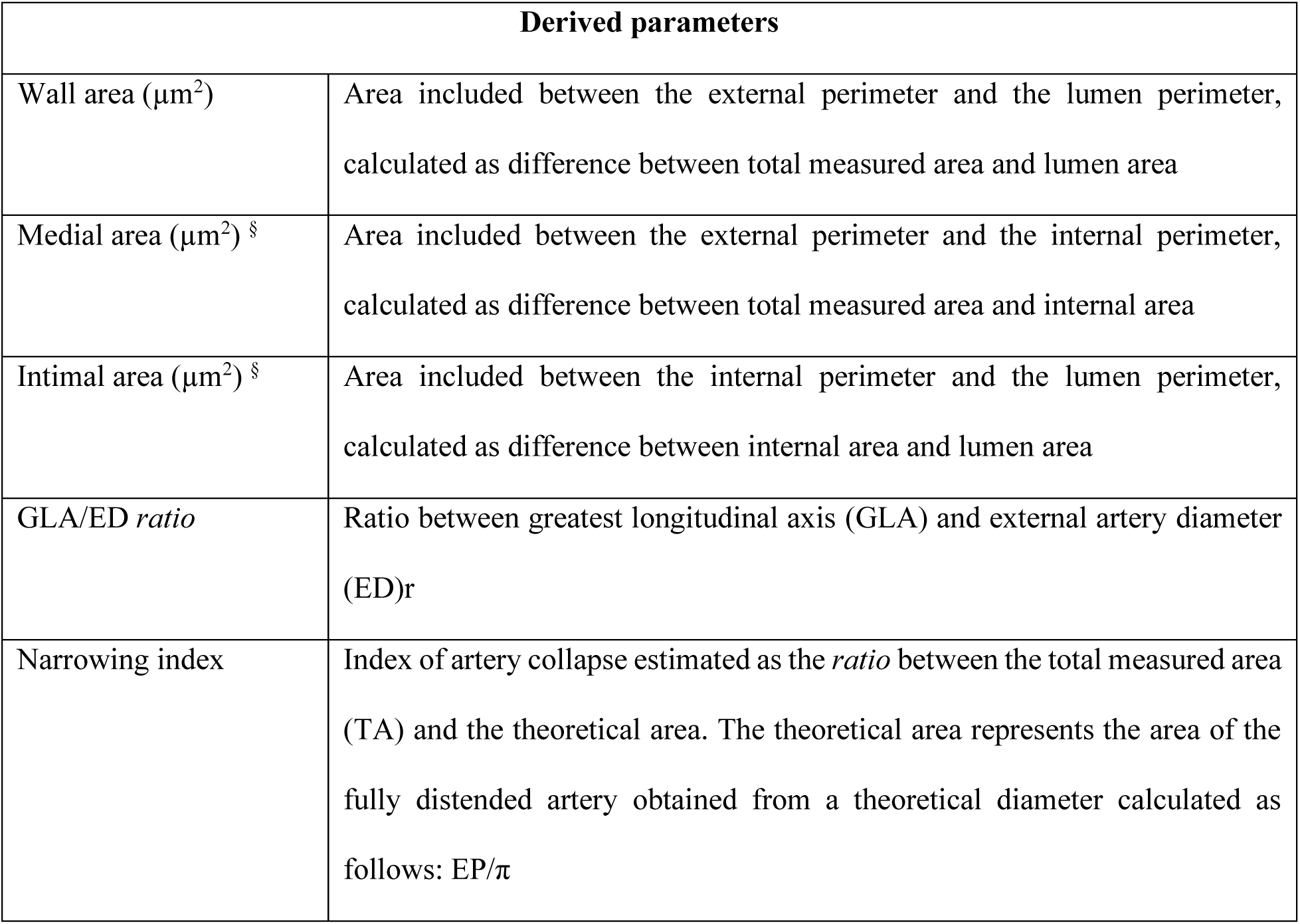
2D derived parameters. These parameters were adapted from those already defined in pulmonary vascular histomorphometric studies in human chronic obstructive pulmonary disease (COPD) (30). Those parameters marked with ^§^ were assessed only on sections stained with Movat Russell pentachrome (study 2).

| Derived parameters |  |
| --- | --- |
| Wall area ($\mu\text{m}^2$) | Area included between the external perimeter and the lumen perimeter, calculated as difference between total measured area and lumen area |
| Medial area ($\mu\text{m}^2$) <sup>§</sup> | Area included between the external perimeter and the internal perimeter, calculated as difference between total measured area and internal area |
| Intimal area ($\mu\text{m}^2$) <sup>§</sup> | Area included between the internal perimeter and the lumen perimeter, calculated as difference between internal area and lumen area |
| GLA/ED *ratio* | Ratio between greatest longitudinal axis (GLA) and external artery diameter (ED)r |
| Narrowing index | Index of artery collapse estimated as the *ratio* between the total measured area (TA) and the theoretical area. The theoretical area represents the area of the fully distended artery obtained from a theoretical diameter calculated as follows: $EP/\pi$ |

**Figure 1:**
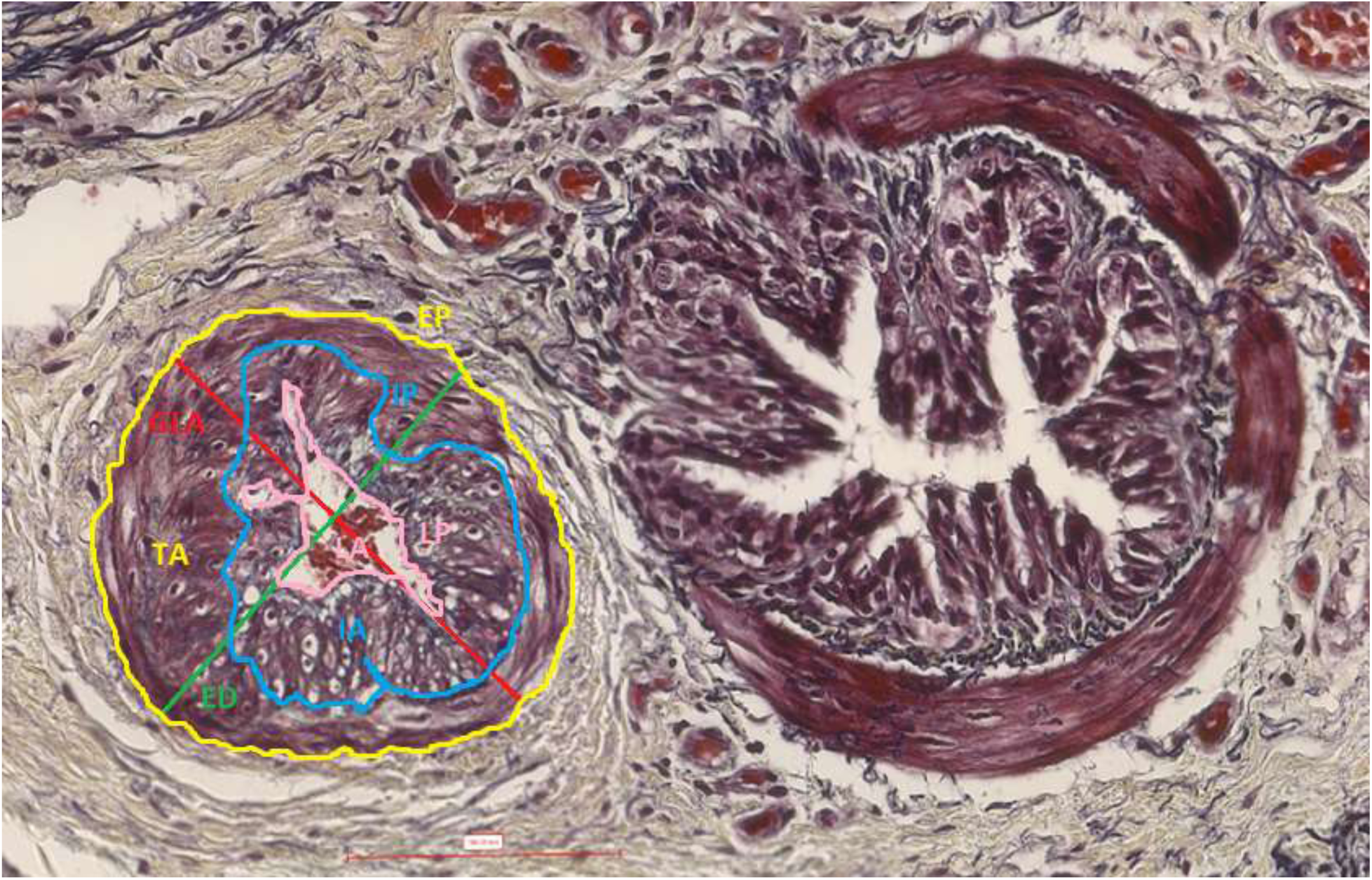
histomorphometric 1D and 2D measured parameters (muscular pulmonary artery and annexed bronchus histological section, 40X, Movat Russel pentachrome). GLA (red outline) = great longitudinal axis; ED (green outline) = external diameter; EP (yellow outline) = external perimeter; IP (blue outline) = internal perimeter; LP (pink outline) = lumen perimeter; TA (within yellow outline) = total area; IA (within blue outline) = internal area; LA (within pink outline) = *lumen* area.

Immunostained muscular pulmonary arteries were scanned at 40X magnification and digitalized as SVS images with PanOptiq™ software version 1.4.3 (ViewsIQ, Vancouver, Canada). Histomorphometry was then performed on the SVS images using the NewCAST™ software version 4.5.1.324 (Visiopharm Integrator System, Hoersholm, Denmark). The total vessel area (µm^2^) and the α-SMA positive and negative wall area (µm^2^) were estimated with a counting point technique. Briefly, on each uploaded SVS digital image, every visible pulmonary artery section was delimited as a region of interest (ROI), using the “ROI drawing” function. The parameters counted were the sum of the points falling on vessel lumen and wall surfaces (∑p_vessel_), the sum of the points falling on α-SMA positive wall surface (∑p_α-SMA+_) and the sum of the points falling on α-SMA negative wall surface (∑p_α-SMA-_). A grid of 576 crosses per screen was applied providing a total count of *at least* 400 points for every counted parameter in each horse ensuring reliable individual estimation according to the current stereology guidelines (32). The total vessel area, the α-SMA positive wall area and the α-SMA negative wall area were estimated by multiplying the ∑p_vessel_, ∑p_α-SMA+_ or ∑p_α-SMA-_ respectively per the surface subtended to a single point (117,22 µm^2^). The total vessel area reflects the overall surface occupied by the artery section (both *lumen* and wall surface), while the α-SMA positive wall area and the α-SMA negative area reflect respectively the PASM and ECM mass within the artery wall (figure 2). Both α-SMA positive and negative wall areas were then expressed as a percentage of the total vessel area (α-SMA + area %; α-SMA - area %), to allow comparison among data of different sized-artery sections.

**Figure 2:**
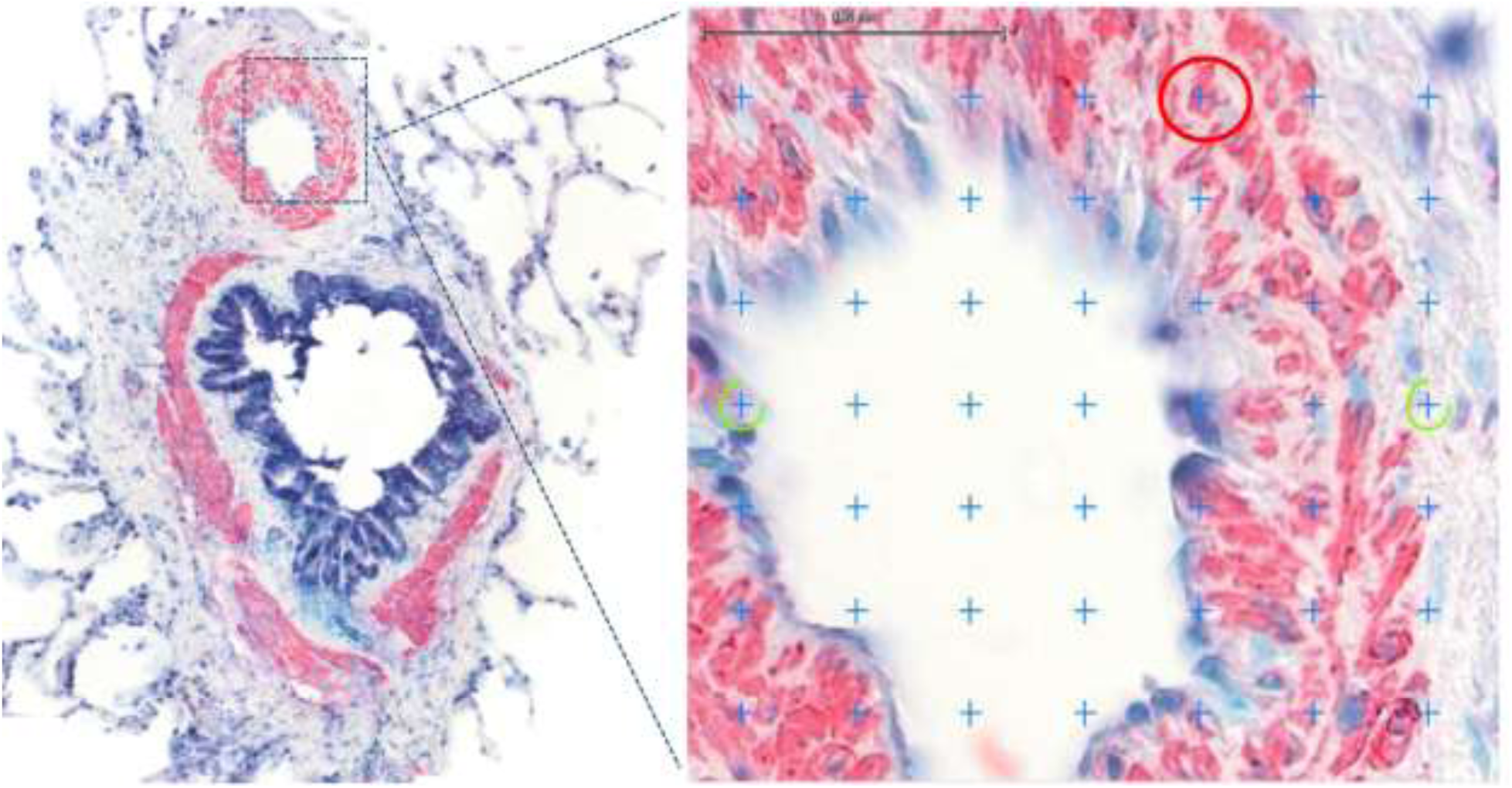
immunostaining for α-SMA and point counting analysis. Pulmonary arteries smooth muscle (PASM) stained pink-red with immunostaining for α-smooth muscle actin (α-SMA), while extracellular matrix (ECM) did not stain. An example of region of interest (ROI) analysis with the point counting technique is provided. A grid of 576 crosses/screen was applied. Crosses falling on pink-red artery wall points (red circle) were counted as α-SMA+ (representative of PASM). Crosses falling on uncolored artery wall points (yellow circle) were counted as α-SMA-(representative of ECM).

### Statistical analysis

Previous studies have demonstrated statistically significant differences in lung tissue remodeling and its reversibility between asthmatic horses and controls using samples of 5-6 horses in each group (22, 26). Therefore, samples from 6 different horses for each group (controls, asthma remission and asthma exacerbation) were included to perform the preliminary study (study 1). Preliminary data obtained in study 1 were used to perform a power analysis, confirming that tissue samples from 5 horses per group will be sufficient to demonstrate statistically significant differences between controls and asthmatic horses in remission (α = 0.05; *p* = 0.90). Because groups were small, normality was assessed by visual inspection of the data. All comparisons were performed using *t* Student test with Welch’s correction (study 1 and 2) and paired *t* test (study 3). One-tailed tests were used to increase power in comparing groups for the wall area%, intimal area%, medial area%, α-SMA+ and - area %, because according to the hypothesis, significant changes were expected only in one direction (increase in asthmatic samples for study 1 and 2, decrease at follow up for study 3). Statistical tests were performed with GraphPad Prism™ software v. 6 (GraphPad software Inc., California, USA). The level of statistical significance was set at p < 0.05.

## Results

Table 3 summarizes the average numbers of histological and immunostained arteries assessed in each study.

**Table 3:** total number, median and ranges of the histological and immunostained pulmonary arteries assessed. ^§^In study 3, one horse of the antigen avoidance group was not considered for the statistical analysis of the pulmonary arteries smooth muscle mass quantification because at the baseline only one immunostained artery was measurable.

| Study /Group | Horses<br>(n) | Histology |  |  | Immunostaining |  |  |
| --- | --- | --- | --- | --- | --- | --- | --- |
|  |  | Arteries<br>(total) | Median/<br>horse | Range | Arteries<br>(total) | Median/<br>horse | Range |
| Study 1 |  |  |  |  |  |  |  |
| Controls | 6 | 222 | 37 | 19 - 51 | - | - | - |
| Asthma | 12 | 502 | 36 | 27 - 68 | - | - | - |
| - Remission | 6 | 272 | 43 | 31-66 |  |  |  |
| - Exacerbation | 6 | 230 | 33 | 27 - 68 |  |  |  |
| Study 2 |  |  |  |  |  |  |  |
| Controls | 5 | 93 | 15 | 10 - 32 | 111 | 22 | 14 - 32 |
| Asthma | 6 | 116 | 17 | 7 - 44 | 114 | 17 | 8 - 33 |
| Study 3 |  |  |  |  |  |  |  |
| Antigen avoidance |  |  |  |  |  |  |  |
| - Baseline | 5 (4) § | 58 | 7 | 2 - 26 | 73 § | 15 § | 11 – 32 § |
| - Follow up | 5 (4) § | 126 | 26 | 4 - 42 | 51 § | 13 § | 6 – 19 § |
| Corticosteroids |  |  |  |  |  |  |  |
| - Baseline | 6 | 46 | 8 | 4 - 12 | 74 | 12 | 4 -23 |
| - Follow up | 6 | 123 | 18 | 13 - 36 | 97 | 14 | 6 - 32 |

### Study 1

Figures 3 and 4 summarize results from study 1. Briefly, asthmatic horses had significantly less collapsed (greater narrowing index) and more thickened (greater wall area %) pulmonary arteries, compared to controls. However, wall area % was similar in severe asthmatic horses during exacerbation and remission. Pulmonary artery wall thickening (greater wall area %) was detected in the apex) and caudodorsal lung fields, but not in the main lobe. Significant increase in wall area % was also detected in the specific region sampled via thoracoscopy.

**Figure 3:**
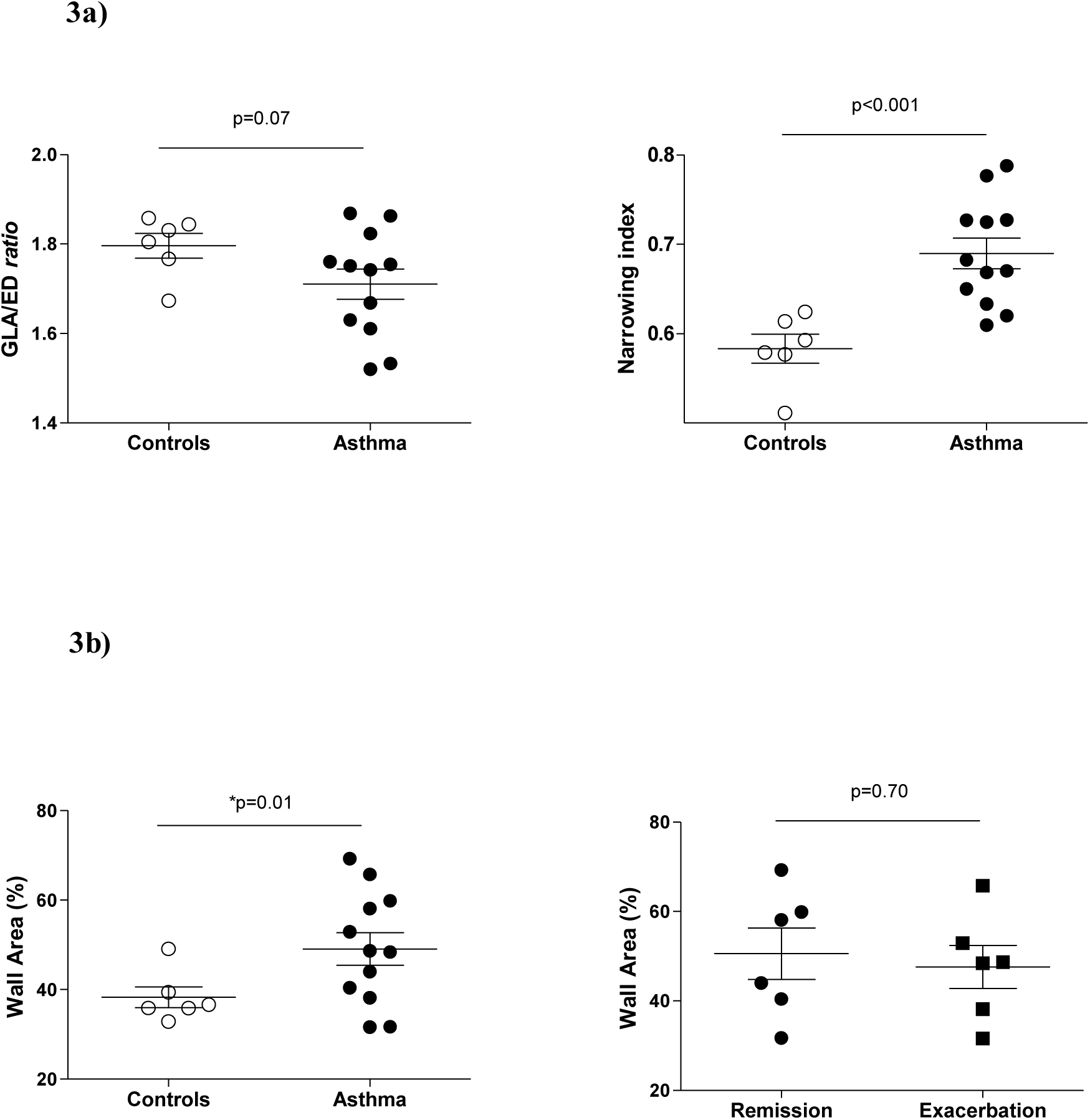
summary of study 1 results. Figure 3a: GLA (great longitudinal axis)/ED (external diameter) *ratio* and narrowing index. Asthmatic horses and controls did not differ in respect to histologic cut sections (GLA/ED *ratio*) but a significantly higher narrowing index was detected in asthmatic horses, indicating lower degree of *post-mortem* vascular collapse. **Figure 3b: wall area %**. Asthmatic horses showed significant wall thickening (increased wall area %) compared to controls, however no difference was detected between remission and exacerbation.

**Figure 4:**
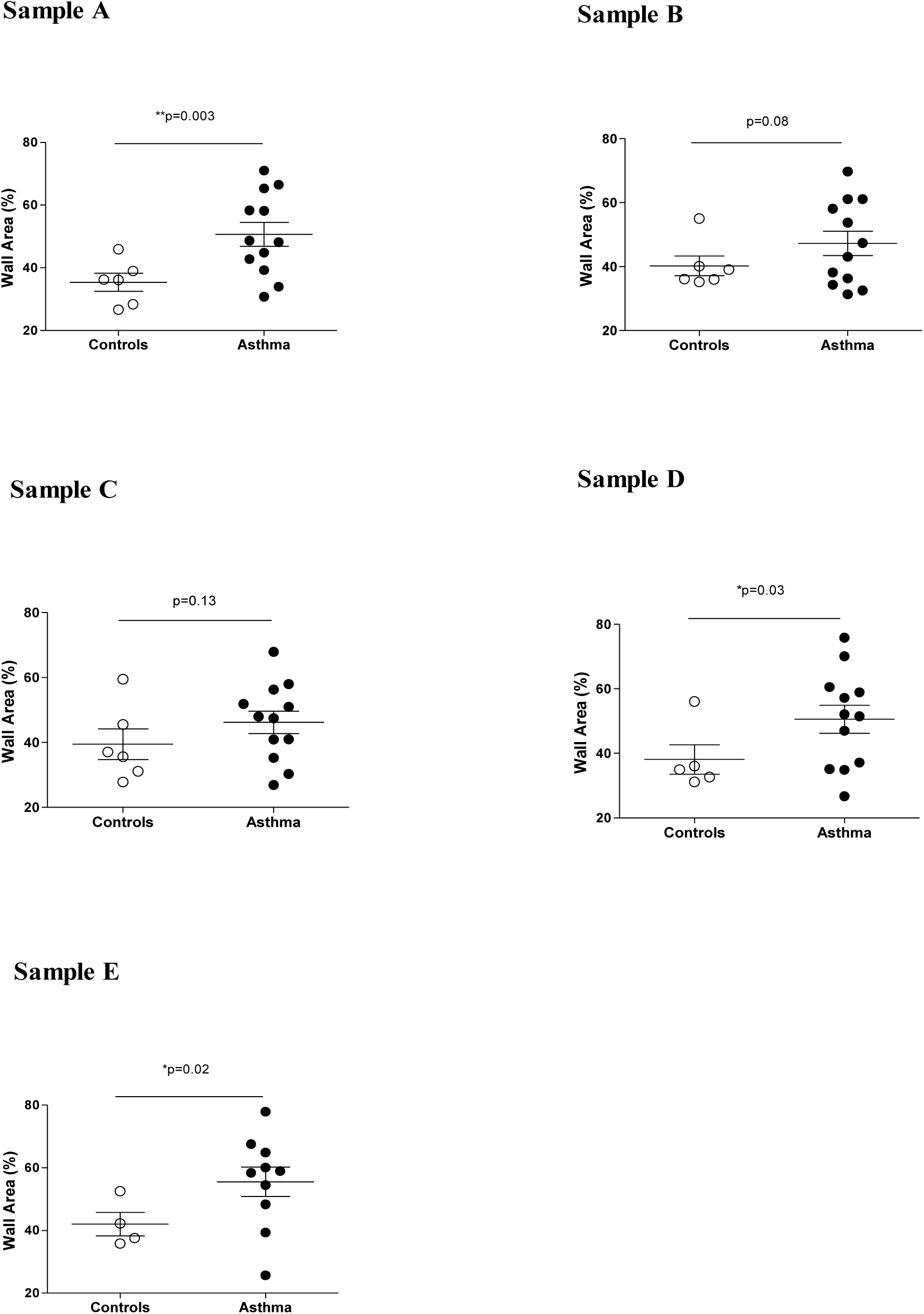
summary of study 1 results: wall area % in different lung regions (samples A-E). Significant wall thickening (increased wall area %) was detected in sample A, sample D and sample E. As previously described (1), sample A was collected from the lung apex, sample B at the emergence of the main bronchus, sample C from the center of the main lung lobe, sample D from the caudodorsal part of the main lung lobe and sample E was collected from the peripheral caudodorsal part of the main lung lobe and corresponds to the specific biopsy site during thoracoscopy.

### Study 2

Figure 5 and figure 6 summarize results from study 2. Conversely to study 1, asthmatic horses and controls had similar degree of collapse (similar narrowing index). Similarly to study 1, asthmatic horses showed significant wall thickening (greater wall area %) compared to controls, however, it was not associated with a specific increase in % of medial area or % of intimal area. Immunostaining revealed that in asthmatic horses, the increase in % of wall area corresponded to a significant increase in the % of PASM mass but not in ECM.

**Figure 5:**
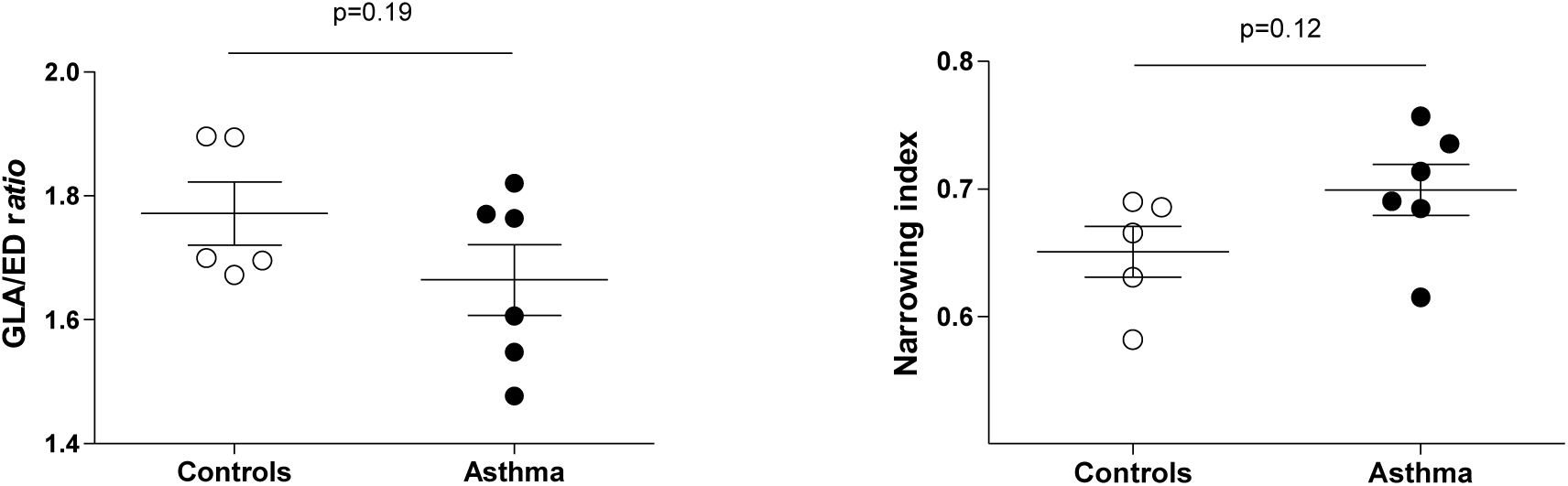
summary of study 2 results: GLA (great longitudinal axis)/ED (external diameter) *ratio* and narrowing index. Asthmatic horses and controls did not differ in respect to histologic cut sections (GLA/ED *ratio*) and degree of vascular collapse (narrowing index).

**Figure 6:**
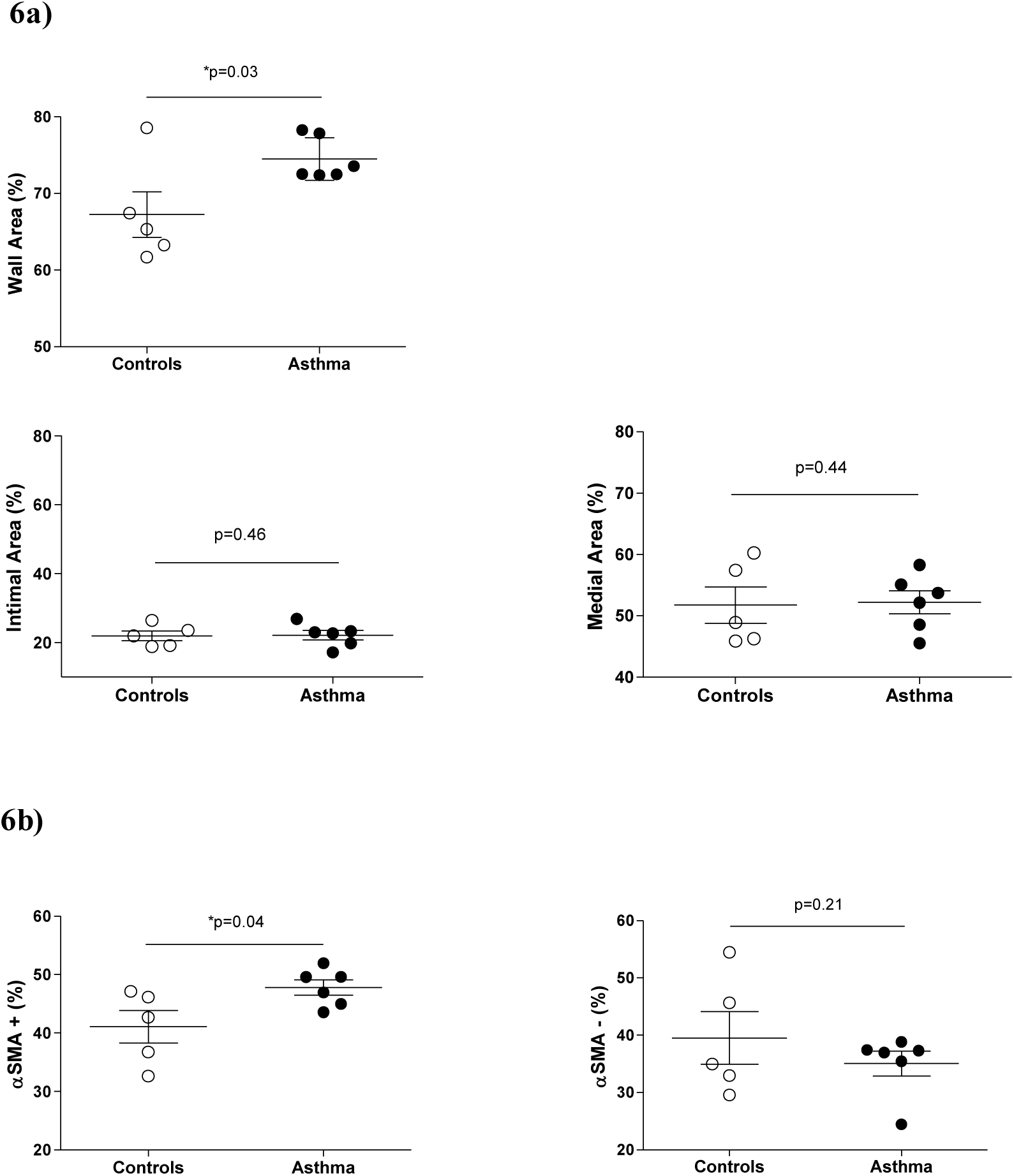
summary of study 2 results. Figure 6a: wall area %. Asthmatic horses showed significant wall thickening (increased wall area %) compared to controls, however no specific increase in medial or intimal area was detected. **Figure 6b: α-SMA+ (α-smooth muscle actin positive) area % and α-SMA- (α-smooth muscle actin negative area) %**. Asthmatic horses showed significant increase in the pulmonary arteries smooth muscle mass (PASM; α-SMA+ area %) but not in the extracellular matrix (ECM; α-SMA-area).

**Figure 7:**
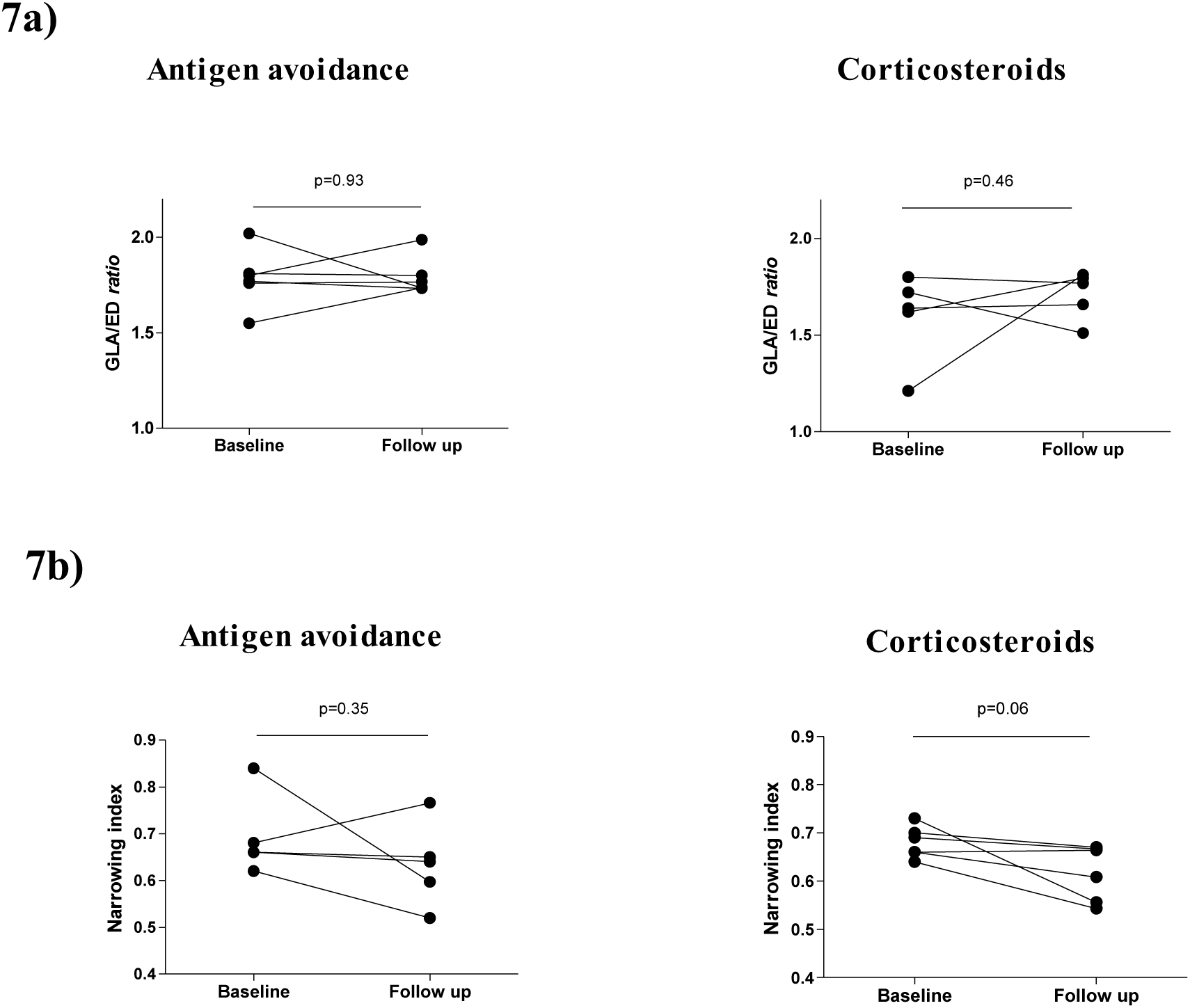
summary of study 3 results. Figure 7a: GLA (great longitudinal axis)/ED (external diameter) *ratio*. In both treatment groups, no differences in respect to histologic cut sections (GLA/ED *ratio*) were found between baseline and follow up. **Figure 7b**: **narrowing index**. No differences between baseline and follow-up were detected in the antigen avoidance group. In the corticosteroid treated group, narrowing index showed a tendency to be reduced at the follow-up.

**Figure 8:**
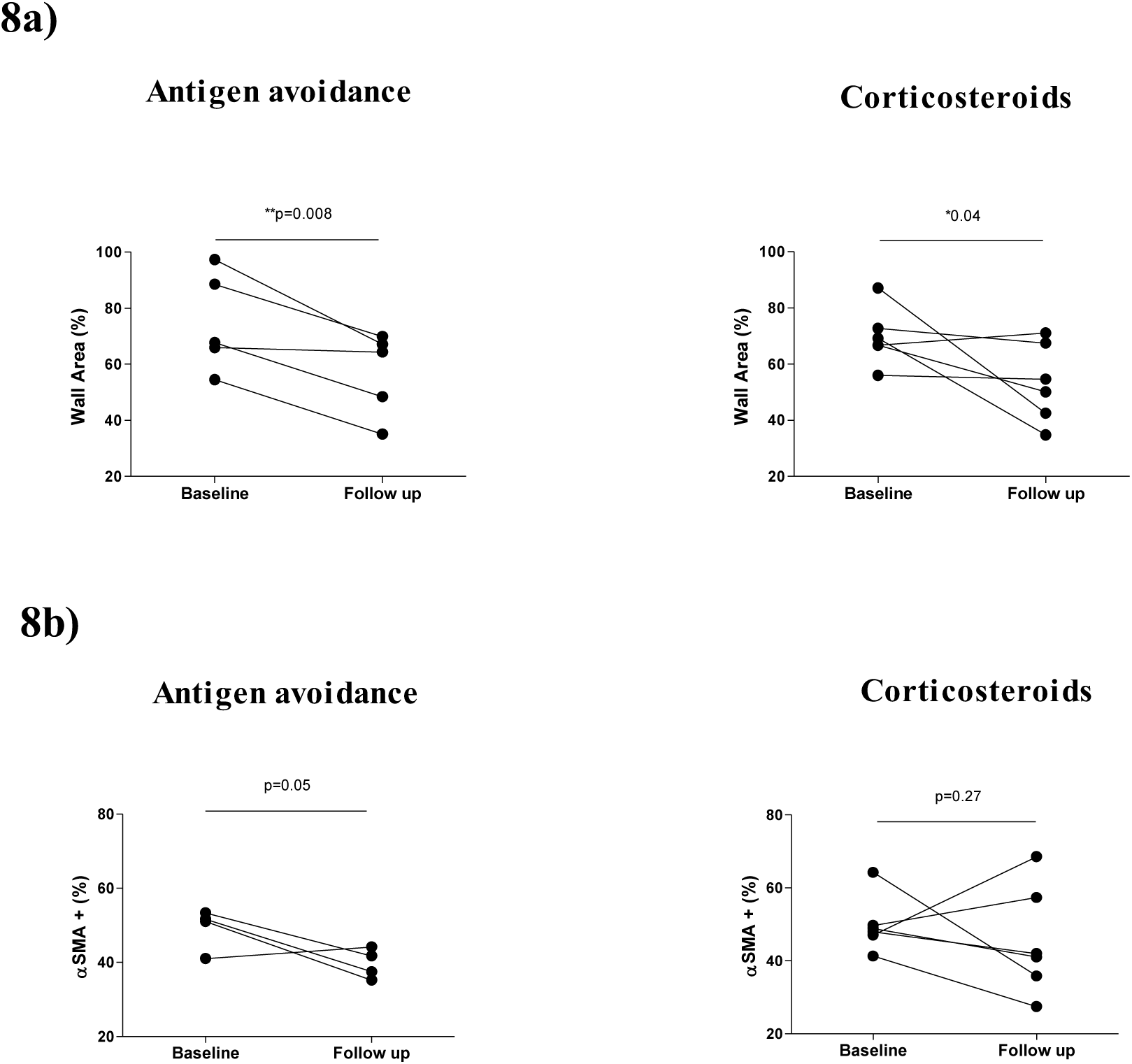
summary of study 3 results Figure 8a: wall area %. In both treatment groups, a significant reduction of the wall thickening (wall area%) was observed at the follow-up, although reduction was more uniform in the antigen avoidance group. **Figure 8b: α-SMA+ (α-smooth muscle actin positive) area %**. Although not significant, tendency to uniform reduction in pulmonary arteries smooth muscle mass (PASM; α-SMA+ area %) at the follow-up was observed in the antigen avoidance group but not in the corticosteroid treated group.

### Study 3

Figure 6 summarizes results from study 3. The narrowing index (degree of collapse) showed a trend towards decreasing in the corticosteroid treated group while it was unaffected in the antigen avoidance group. In both groups, long-term treatment induced significant reduction of the wall area. However, only the antigen avoidance strategy was associated with an almost significant PASM % reduction.

## Discussion

The present study revealed significant pulmonary artery wall thickening (increase in the pulmonary artery wall area) in asthmatic horses compared to controls. Remodeling involved the lung apex and the caudodorsal lung fields, including the specific region sampled by *in viv*o thoracoscopy. In asthmatic horses, pulmonary artery wall thickening was due to an increase in PASM mass, but the ECM was unaffected. Remodeling changes (wall area and PASM increase) were uniformly reversible after long-term antigen avoidance (12 months), but not after 2 to 4 months, neither after 12 months of inhaled corticosteroids combined with 6 months of antigen avoidance.

This is the first study that used a quantitative histomorphometric approach to assess pulmonary vascular remodeling in horses. The mean wall area of muscular pulmonary arteries in asthmatic horses was about 7% and 12% greater than that of controls, *in vivo* and *post-mortem* respectively. Increases of similar magnitudes (7-10%) were observed in the lungs of COPD-affected human patients compared to controls (30, 33, 34). *Post-mortem*, pulmonary arteries of asthmatic horses appeared significantly less collapsed (higher narrowing index) than those of controls. This difference was not observed *in vivo*, with both groups showing similar low degree of collapse. Pulmonary arteries of asthmatic horses may show hypercontractility and impaired *post-mortem* relaxation similarly to what previously demonstrated in experimental animal models of lung allergic inflammation (35-37). However, differences in wall area (expression of wall thickening) were not affected by reactivity differences between groups and were similarly detected both *in vivo* and *post-mortem*, suggesting consistent remodeling of pulmonary artery walls in asthmatic horses.

Vascular remodeling affected only the caudodorsal lung fields and the lung apex. Lung blood flow distribution in horses increases in a dorsal and caudal direction (38, 39). Higher perfusion of caudodorsal lungs may mean higher delivery of inflammatory cells and factors during acute lung inflammation, potentially contributing to selectively located remodeling. Caudodorsal lung fields represented the main site of vascular remodeling also in equine exercise-induced pulmonary hemorrhage (EIPH) (38, 40). Presence of vascular remodeling in the lung apex was surprising, however. Despite low blood flow, cranioventral small pulmonary arteries appeared more prone to remodeling also in a previous study (41). The causes are likely due to regional differences in vessel mechanical reactions in response to yet unidentified stimuli that may also be evoked in severe equine asthma. As a hypothesis, endothelial dysfunction may occur in severe equine asthma as described in human COPD. This resulted in impaired production of vasodilators, particularly nitric oxide which vasodilating role was significant mainly in the cranioventral lung (20, 33, 34, 42). Of note, pulmonary artery remodeling was demonstrated *post-mortem* in the same region sampled by *in vivo* thoracoscopy (sample E). This technique appeared adequate to investigate prospectively *in vivo* vascular remodeling. Independently from the group, pulmonary arteries were 25-30% thicker when lung samples were collected *in vivo* than *post-mortem*. Methodological factors may have been the cause of the difference. In humans, minimally invasive techniques applying high traction force to harvest systemic vessels impair endothelium-dependent relaxation, without affecting contractile response (43).

Presence of vascular remodeling in asthmatic horses was supported by the significant increase in the PASM, in agreement with what reported in other chronic spontaneous lung disorders, such as human COPD or equine EIPH, and in rodent asthma models (17-19, 40, 44-46). In EIPH, a common condition in high intensity performing horses, pulmonary arteries remodeling was also characterized by artery wall thickening with medial smooth muscle hypertrophy, intimal hyperplasia, elastic lamina inconsistency and lack of organization (40). Severe asthmatic horses did not show specific thickening of the intimal and/or medial layer, however. Pulmonary artery remodeling could have been irregular with total increase of wall PASM mass that was not consistent within a single layer. Absence of specific intimal/medial thickening could also depend on the presence of some degree of elastic lamina inconsistency that prevented intimal and medial area differentiation in all arteries and reduced power of the histomorphometric assessment. Remodeling itself may have caused loss of internal elastic lamina integrity and staining definition, as occurring in equine EIPH.

Persistent hypoxic pulmonary vasoconstriction due to chronic hypoxia resulted in PASM “work” hypertrophy in experimental animal models and a similar pathophysiology could explain the PASM increase of severe asthmatic horses (9, 10, 47). Airway inflammation may also have contributed. In rodent models, allergic airway inflammation increased PASM mass and cell turnover with similar time course and magnitude to airway smooth muscle remodeling, both being not reversible after one month of antigen avoidance (17, 18). An increase in airway smooth muscle mass was also present in severe asthmatic horses (22, 26). Short-term antigen avoidance significantly improved lung function and airway inflammation in asthmatic horses, while smooth muscle mass was not reduced in muscular pulmonary arteries in the present study, nor in small peripheral airways in a previous work (22). As the PASM cell turnover was not investigated, whether the increase in PASM mass was due to cell proliferation, hyperplasia or hypertrophy was not determined.

In severe asthmatic horses, the increase in pulmonary artery PASM mass could contribute to pulmonary hypertension during disease exacerbation, by enhancing magnitude of hypoxic pulmonary vasoconstriction. Experimentally induced hypoxemia increases mean pulmonary artery pressure both in normal and in asthmatic horses; however, only asthmatic horses develop pulmonary hypertension (12). Furthermore, persistence of PASM mass increase and related artery wall thickening may explain the significantly higher mean pulmonary artery pressure values, although within physiological ranges, in asthmatic horses during remission than in controls (12). In the present study, pulmonary artery remodeling did not affect the amount of vascular ECM. In rodents, increased collagen was detected during sustained exacerbation mainly in the perivascular layer that was not examined in the present study (17, 19). Perivascular changes and especially ECM deposition merit to be further investigated in severe equine asthma.

Pulmonary artery remodeling (wall thickening and PASM increase) was reversed by long-term (12 months) antigen avoidance. In horses treated with long-term (12 months) inhaled corticosteroids, remodeling could not be effectively reversed, in spite of 6 months of concomitant antigen avoidance. While inhaled corticosteroids were mainly effective on airway obstruction, long-term antigen avoidance better controlled neutrophilic airway inflammation (24). This supported a possible role of inflammation in the development of pulmonary artery remodeling, facilitated by the strict anatomic relationship between pulmonary circulation and the airways, as already investigated in acute allergic human asthma (48). Otherwise, treatment with inhaled corticosteroids resulted in significant vasorelaxation, as expressed by the higher degree of collapse (lower narrowing index). Both *in vitro* and experimental *in vivo* studies have shown that corticosteroids may improve the relaxation response of pulmonary arteries by decreasing oxidative stress, up-regulating the expression of nitric oxide synthases and increasing bioavailability of nitric oxide in pulmonary endothelial cells (49, 50).

In conclusion, the presence of pulmonary artery remodeling represented a new finding in equine asthma. It was possible to study the reversibility of PASM mass increase, achieved only with long-term antigen avoidance strategies, suggesting a role for inflammation in the persistence of the vascular changes. Mechanisms of PASM remodeling, endothelium and perivascular changes remains to be investigated as well as the possible alterations of distal arterioles, pulmonary capillaries and veins. Further studies should focus on the role of hypoxemia and inflammation in inducing PASM remodeling as well on its impact on pulmonary artery pressure and cardiovascular complications in severe equine asthma.

## Acknowledgments

The authors would like to acknowledge Amandine Vargas for her support in tissue processing, staining and imaging and Guy Beauchamp for his support with statistical analysis.

